# Approaching object acceleration differentially affects the predictions of neuronal collision-avoidance models

**DOI:** 10.1101/2022.10.26.513898

**Authors:** Fabrizio Gabbiani, Thomas Preuss, Richard B. Dewell

## Abstract

The processing of visual information for collision avoidance has been investigated at the biophysical level in several model systems. In grasshoppers, the *η* model captures reasonably well the visual processing performed by an identified neuron called the lobular giant movement detector (LGMD) as it tracks approaching objects. Similar phenomenological models have been used to describe either the firing rate or the membrane potential of neurons responsible for visually-guided collision avoidance in other animals. In goldfish, the *κ* model has been proposed to describe the Mauthner cell, an identified neuron involved in startle escape responses. In the vinegar fly, a third model was developed for the giant fiber neuron, which trigger last resort escapes immediately before an impending collision. One key property of these models is their prediction that peak neuronal responses occur a fixed delay after the simulated approaching object reaches a threshold angular size on the retina. This prediction is valid for simulated objects approaching at a constant speed. We tested whether it remains valid when approaching objects accelerate. After characterizing and comparing the models’ responses to accelerating and constant speed stimuli, we find that the prediction holds true for the *κ* and the giant fiber model, but not for the *η* model. These results suggest that acceleration in the approach trajectory of an object may help distinguish and further constrain the neuronal computations required for collision avoidance in grasshoppers, fish, and vinegar flies.

## 1 Introduction

Visually guided collision avoidance has proven attractive to study how the nervous system processes visual information to generate escape behavior and to investigate the cognitive processes implicated in the various escape strategies adopted by prey [9]. At the biophysical level, several models have guided the study of collision-detecting neurons, helping to shed light on the coding and processing of visual information by single neurons [12, 21]. Three distinct such models were designed to fit data in grasshoppers, vinegar flies and goldfish [16, 34, 1, 24]. These models were fitted to reproduce responses to looming stimuli, that simulate the approach of an object at constant speed towards the potential prey [28]. A common feature of these models is the prediction that neuronal responses peak at a fixed delay before the approaching object reaches a threshold angular size on the animal’s retina [15]. Here, we ask whether the models make distinct predictions for peak neural responses when presented with different stimuli that simulate an approach with an acceleration different from zero. Although the impact of acceleration on the responses of collision-detecting neurons has not been tested in a controlled setting, it is known to be ethologically relevant in at least one case: the predation of vinegar flies by damselflies [33]. Our results suggest that accelerating stimuli could shed further light on the biophysical processing of visual information for collision detection and that they may help distinguish and refine models of visually guided collision avoidance experimentally. In a companion manuscript, we use the new stimuli and model predictions to gain further insight in the neural computations underlying collision-avoidance behaviors of grasshoppers and goldfish. As more is learned about the approach trajectories of predators, these models may also help explain why specific predation strategies are more successful than others.

The remaining of this paper is organized as follows: after a brief explanation on methods and notation (sec. 2), we introduce looming stimuli (sec. 3) and then the models describing the response of collision-detecting neurons to such stimuli (sec. 4). In section 4, we also derive the main properties of the models’ response to looming stimuli, illustrating their similarities and differences, and we show that the models can be mapped onto each other under an assumption that preserves their main feature and that is required to describe the response variability of collision-detecting neurons across grasshoppers. In sec. 5, we introduce stimuli approaching with a constant acceleration (or deceleration) and show how the various neuron models differ in their responses to such stimuli. Sec. 6 introduces a second set of stimuli approaching with a time-dependent deceleration and shows how they can be used to distinguish the different models. We conclude by a brief summary and discussion of the results (sec. 7).

## 2 Methods

For the *η* and *κ* models algebraic results could be derived by hand and were verified through Matlab simulations. For the giant fiber model, the predictions were tested using Matlab only as the model structure precludes pen-and-paper derivations.

## 3 Looming Stimuli

Classically, looming stimuli have been defined as the simulation on a two-dimensional screen of objects approaching at a constant speed on a collision course with an animal [28]. For a solid square of half-size *l*, starting at an initial distance, *x_i_*, the distance from the eye as a function of time *s* ≥ 0 measured from movement onset is given by:

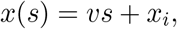

where *v* < 0 is the approach speed. The angular size subtended by the object at the eye depends on the ratio of the object’s distance to its half-size. Hence, it will be useful to define the normalized distance *y*(*s*) = *x*(*s*)/*l* so that

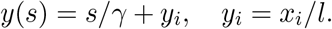

The constant *γ* = *l/v* < 0 (in units of time) fully characterizes the approach trajectory. Solving *y*(*s*) = 0 yields the time of collision, *s_c_* = –*γy_i_* > 0 (Fig. 1**A**). If time is referenced relative to collision, *t* = *s* – *s_c_* ≤ 0, then *y*(*t*) = *t/γ* (Fig. 1**B**). The half-angle subtended by the object at the retina is then obtained by trigonometry:

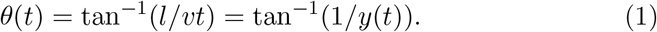

It is an expanding function of *y*(*t*) that grows increasingly fast towards collision time (Fig. 1**C**). The corresponding angular velocity, also increases non-linearly, but eventually saturates towards collision time (Fig. 1**D**). In several species, typical values for the parameter *γ* leading to successful escape behaviors range from −80 to −20 ms (e.g., [24, 11, 36]). In grasshoppers and locusts, this range extends further, reaching −120 ms [10]. For the same species, a typical initial angular half-size *θ_i_* = tan^−1^(1/*y_i_*) of ~ 0.75° will be below the spatial resolution of the eye (~ 2°) and minimize transient onset neuronal responses at the start of the approach.

**Figure 1:**
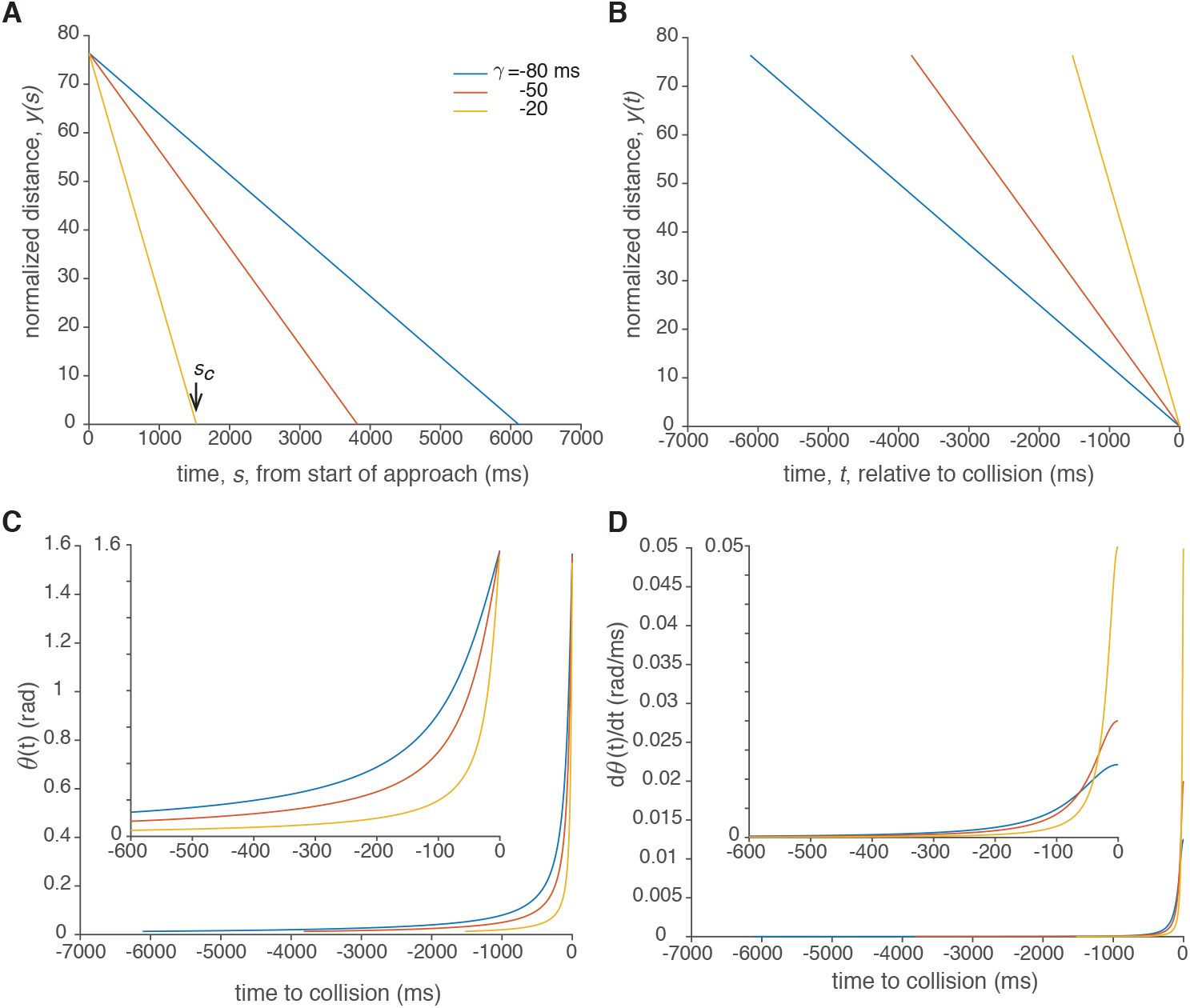
**A**, Normalized distance of the approaching object, *y*(*s*), as a function of time from approach onset for three *γ* = *l/v* values (−80, −50 and −20 ms). The initial value, *y_i_* = 76.4, corresponds to an initial halfangle of 0.75°. The vertical arrow indicates collision time, *s_c_*, for the fastest approach trajectory. **B**, Same three approaches with time referenced relative to collision (*y*(*t* = 0) = 0). **C**, Corresponding half-angle, *θ(t),* subtended by the object at the retina. Collision occurs when *θ* = 90 degrees or 1.57 radians. **D**, Corresponding angular velocity, 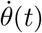, during approach. In **C**, **D**, the *inset* magnifies the last 600 ms of approach.

## 4 Neural models of collision avoidance

### Eta model

In grasshoppers and locusts, a pair of identified neurons is chiefly responsible for jump and flight escape behavior in response to looming stimuli. The first of these neurons, the lobula giant movement detector (LGMD), integrates excitatory, motion sensitive inputs and inhibitory, size-dependent inputs across one visual hemifield [20, 16, 26]. Its postsynaptic target, the descending contralateral movement detector (DCMD) neuron, relays the LGMD spiking activity to motor centers generating jump and flight motor output [19, 12]. The DCMD acts as a faithful relay, with each LGMD spike causing one and only one spike in the DCMD [20]. The initial model used to describe the LGMD/DCMD firing rate was the function

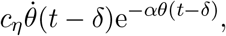

where *c_η_* is a scaling factor converting angular velocity (in rad/s) to firing rate (in spk/s) [16]. This function was later called the ‘*η* function’ [30]. According to this function, the angular velocity acts as an excitatory term since it increases as the object approaches, while the negative exponential acts as an inhibitory term, in agreement with known physiological inputs to the LGMD neuron [26, 14, 23, 37, 35]. As a result, the *η* function predicts an initial increase in firing rate followed by a peak and an eventual decrease when the negative exponential term becomes vanishingly small (Fig. 2**A**). The time of peak, *t_p_*, is determined by the equation *dη/dt* = 0 which leads to

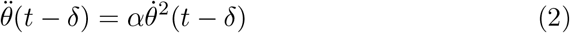

(taking into account that exp(−*αθ*(*t* – *δ*)) ≠ 0). Using eq. 1 yields a linear relation between *t_p_* and the looming stimulus parameter *γ*:

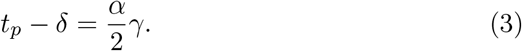

Because the normalized distance is also a linear function of time, with proportionality constant 1/*γ*, the normalized distance at *t_p_* – *δ*, *y*(*t_p_*–*δ*) = *α*/2, is independent of *γ*. Thus, the angle subtended by the stimulus at that time is also independent of *γ*:

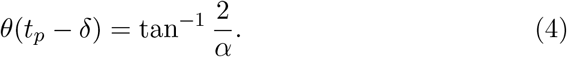

This prediction of a neuronal peak firing rate time occurring a fixed delay (*δ*) after the looming stimulus has reached a threshold angular size has been verified in the LGMD/DCMD neurons [15] and in a number of neurons classes across vertebrate and invertebrate species (e.g., [18, 32, 2, 17, 27, 8]).

**Figure 2:**
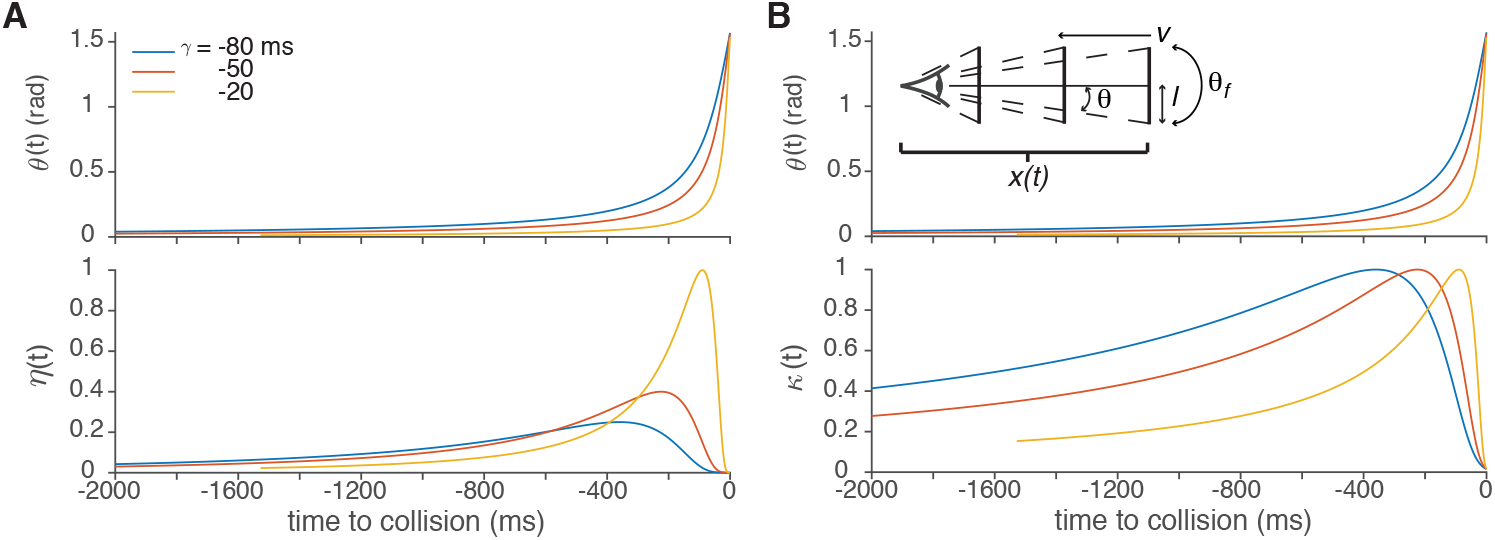
**A**, Angular size (*top*) and responses of the *η* model (*bottom*) to the same three *γ* values as in Fig. 1. The parameter *α* = 9 corresponds to a threshold half-angle of 12.5° (0.22 rad) and *δ* = 0 ms. The peak value for *γ* = −20 ms has been normalized to one. **B**, Corresponding responses of the *κ* model, with the same normalization as in **A**. The parameter *β* = 4.6 was selected to yield the same threshold half-angle as in **A** (see eq. 9). The top *inset* depicts the definition of the kinematical and angular parameters of the stimulus.

### Relation to earlier work

In [15] and other articles, the *η* model was formulated using the full stimulus angle, *θ_f_* = 2 θ, see Fig. 2**B**, inset. Accordingly, the value of the parameter *α* is halved, *α_f_* = *α*/2. Further, the angular edge speed is one half of the full angle expansion speed, 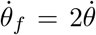. Finally, the stimulus parameter used earlier was *l*/|*v*| = –*γ* (> 0). The parameters selected here facilitate algebraic calculations.

### Additional properties of the eta model

The peak value of the *η* function scales inversely with *γ* since for a looming stimulus,

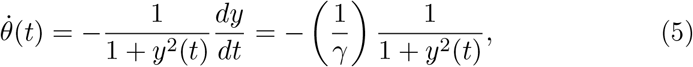

and hence so does the value of *η* at peak time,

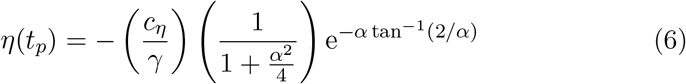

(using eqs.1 and 4, with *δ* = 0; see Fig. 2**A**).

The *η* model also predicts a constant number of spikes prior to collision independent of *γ*. To see this, note first that if *η*(*t*) describes a neuronal firing rate, then the total number of spikes prior to collision is given by

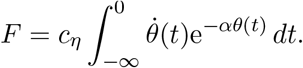

For a looming stimulus, this quantity is independent of the parameter *γ*. Namely, if *γ*_2_ = *γ*_1/*ρ*_, and *y_i_*(*t*) = *t*/*γ_i_*, *i* = 1, 2, then *y*_2_(*t*) = *y*_2_(*ρt*) and consequently,

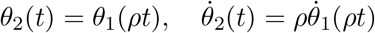

(using eqs.1 and 5). Hence,

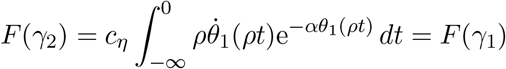

after a change of integration variables, 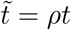.

### Extensions of the eta model

The *η* model works well for a restricted range of *γ* values (in the range [−24; −4] ms; [16]). The model can be naturally extended while preserving the result of eq. 4 by adding to it a static (time-independent) non-linearity, *f*, yielding the composed function *f* (*η*(*t*)) (see [15], appendix 3).

Adequate fits to experimental LGMD/DCMD firing rate data require an additional extension, by making the static non-linearity *γ*-dependent and dependent on whether the fitted firing rate value lies before or after the peak time (see [15], Figs. 12 and 13). This extension leads to the following functional form:

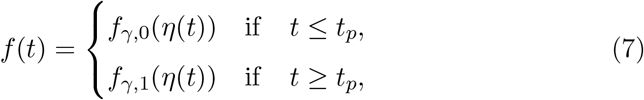

where *t_p_* is given by eq. 3 above. This extended model has been validated over the range *γ* ∈ [−50; −5] ms [15]. With the insight of subsequent work [22, 5, 6, 7], the dependence of the static non-linearity on *γ* and on the rising, resp. decaying, phase of the *η* function is to be expected given the large number of ion channels with activity-dependent kinetics present in the LGMD and shaping its firing rate. These channels will affect differently the rising and decaying phase of LGMD firing due to their time-dependent activation and inactivation kinetics.

### Kappa model

In goldfish, a different model was proposed to describe the time course of the membrane potential in an identified pre-motor neuron, called the Mauthner cell in response to looming stimuli [24]. In this model the membrane potential is a product of the stimulus angular size by a negative exponential of angular size:

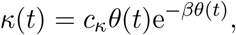

where we have omitted the time delay, *δ*, for simplicity. Just as for the *η* model, the multiplication of *θ* by a negative exponential of *θ* leads to a function that initially increases, then peaks and eventually decays as time to collision nears (Fig. 2**B**). As above, the peak time is found by setting the time derivative of *κ*(*t*) equal to 0 and solving for *t_p_*. Taking into account that 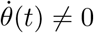 (see Fig. 1**D**) and exp(–*βθ*(*t*)) ≠ 0, we obtain:

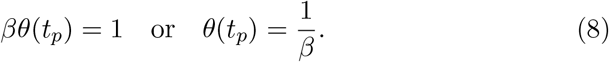

Combining eqs.4 and 8 shows that the *η* and *κ* models predict the same angular threshold size, provided that

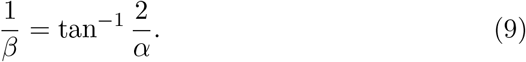

Note that this result remains valid for extensions of the *κ* model by static non-linearities similar to those discussed above for the *η* model. The *κ* model was proposed because it predicts a constant peak value for the membrane potential,

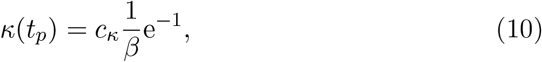

as observed experimentally ([24]; see Fig. 2**B**). In contrast, the *η* function’s peak value scales inversely with *γ* (see eq. 6).

### Equivalence of the eta and kappa models

The *η* and *κ* models are equivalent in the sense that one can be mapped onto the other via a *γ*-dependent and a rising/decaying-phase-dependent static non-linearity, see eq. 7. Specifically, provided eq. 9 holds, the following result transforms the *κ* function into the *η* function for a looming stimulus:

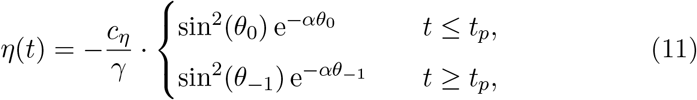

where

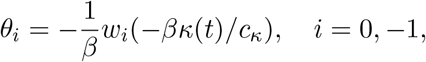

and *w*_0_, *w*_−1_ are the 0^*th*^ and −1^*st*^ branches of the Lambert W function, respectively (Fig. 3).

**Figure 3:**
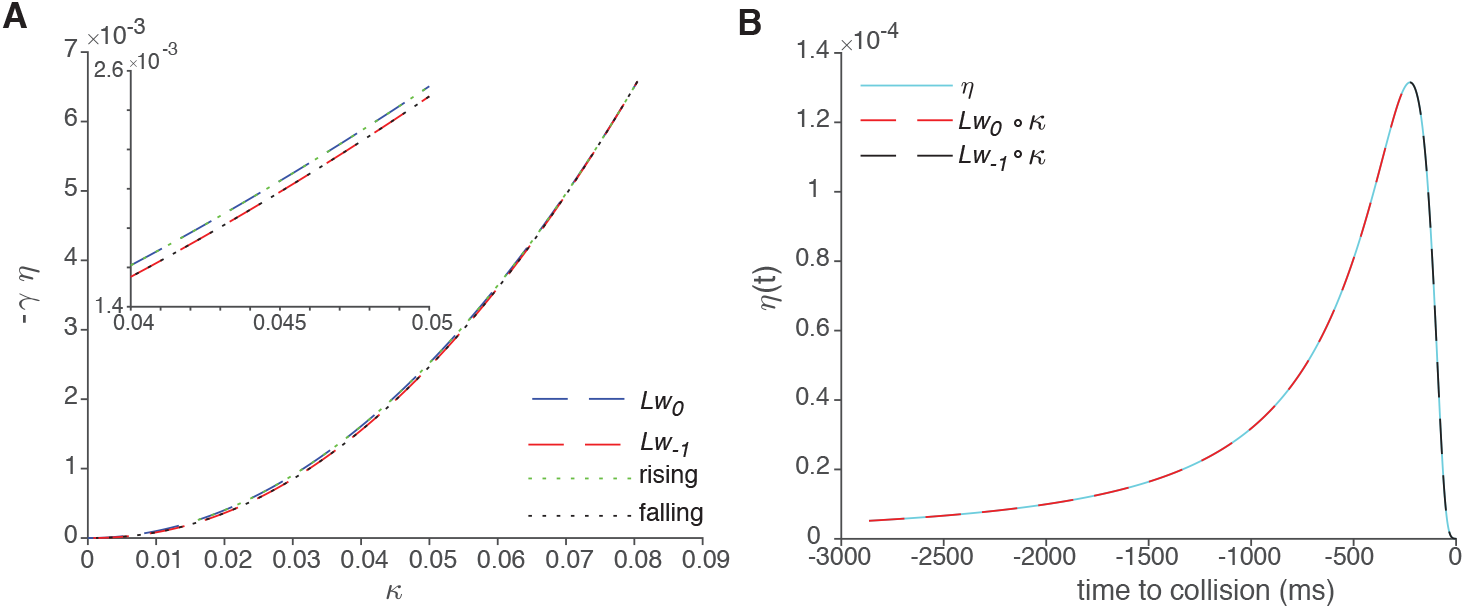
**A**, Nonlinearities of eq. 11 mapping *κ* into *η* and denoted by *Lw*_0_ and *Lw*_−1_, respectively (*Lw_i_*(*κ*) = sin^2^(*θ_i_*(*κ*))e^−*αθ_i_*(*κ*)^, for *i* = 0, −1 and *c_κ_* = *c_η_* = 1). The green and black dotted lines are obtained by mapping numerically *κ*(*t*) onto *η*(*t*) over their rising and falling phases, respectively. The *inset* magnifies the middle portion of the curves. **B**, Time course of *η*(*t*) (cyan, *γ* = −50 ms) and mapping of *κ*(*t*) obtained using eq. 11 (red and black).

**Note**. The Lambert W function is a multivalued function, defined as the inverse of the function *z* → *ze^z^*, where *z* is a complex number. It finds applications in the solution of algebraic equations arising in a variety of scientific fields [4].

**Proof**. We need only consider the case *c_η_* = *c_κ_* = 1. We first use eqs.5 and 1 to express 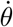 as a function of *θ*:

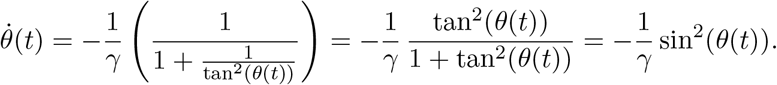

Let now *κ* = *θ*e ^−*βθ*^. We are looking for the solution, *θ* of this equation. First set *z* = −*βθ* or *θ* = –*z/β*. The equation to solve becomes

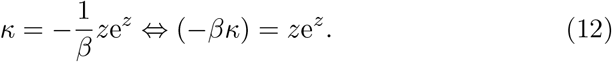

In our case, *κ* lies between 0 and 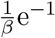 (see eq. 10). Hence, –*βκ* is real and lies between –e^−1^ and 0. Therefore, there are two solutions to eq. 12, namely *w*_0_(–*βκ*) and *w*_−1_(–*βκ*) [4]. The specific assignment of the 0^*th*^ branch to the rising phase of the *κ*(*t*) function is asserted by comparing eq. 11 with its numerical solution (Fig. 3), thus completing the proof.

### Giant Fiber model

This model is based on biophysical experiments that characterized four neuronal inputs to the giant fiber (GF) of *Drosophila melanogaster,* an identified neuron involved in collision avoidance and escape behaviors. The specifics of the model are briefly described here for completness (further details can be found in [1, 34]). It is formulated in terms of the full stimulus angle and angular speed (i.e., 2*θ* and 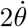 in the notation used here) and relies on the weighted sum of two excitatory and two inhibitory inputs. Thus, the GF membrane potential is modeled

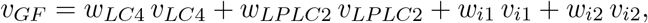

as with weights *w*_*LC*4_ = 1.62, *w*_*LPLC*2_ = 1.45, *w*_*i*1_ = 2.27, and *w*_*i*2_ = 1. The membrane potential of the LC4 neuron encodes the (full) angular velocity:

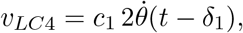

with *c*_1_ = 0.2567 · 10^-3^ mV/(deg/s), and *δ*_1_ = 19 ms. The membrane potential of the LPLC2 neuron model is tuned as a Gaussian for a specific logarithm of the full angular size:

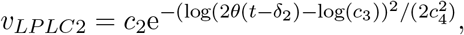

with *c*_2_ = 1.7 mV, *c*_3_ = 42 deg, *c*_4_ = 0.52 (dimensionless) and the delay δ_2_ = 19 ms. The first inhibitory input produces increased inhibition as the stimulus full angular size increases:

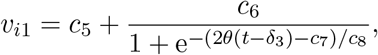

with *c*_5_ = −0.53 mV, *c*_6_ = 0.59 mV, *c*_7_ = 66 deg, *c*_8_ = −11 deg and *δ*_3_ = 37.5 ms. The second inhibitory input is also tuned as a Gaussian for a specific full angular size:

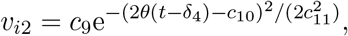

with *c*_9_ = −0.52 mV, *c*_10_ = 26 deg, *c*_11_ = 7.8 deg, and *δ*_4_ = 11 ms. This composite model produces responses similar to those of the *η* and *κ* model (Fig. 4). As in the *η* model, the peak response depends on the parameter *γ*, though not as strongly (compare Fig. **2A** and Fig 4).

**Figure 4:**
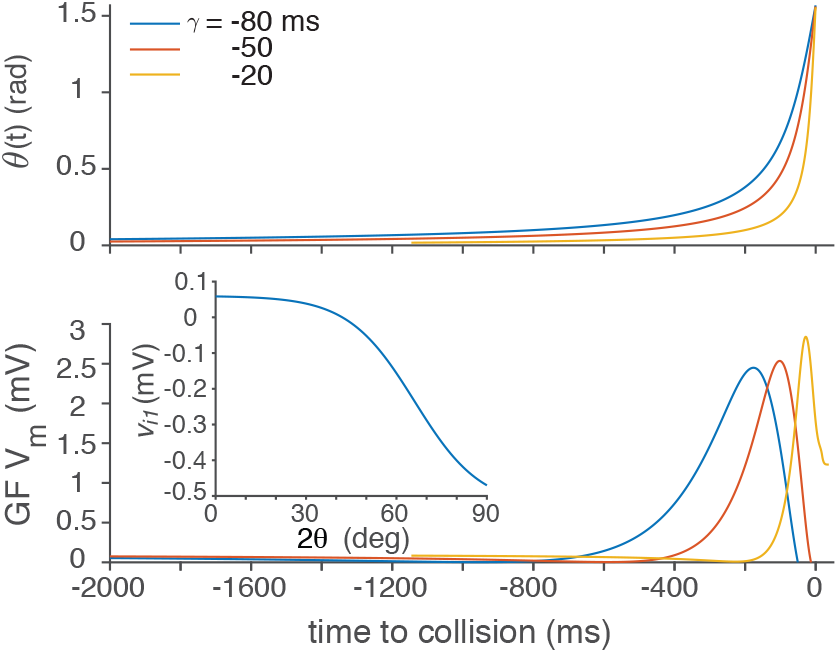
Angular size (*top*) and responses of the GF model (*bottom*) to the same three *γ* values as in Fig. 1. The *bottom inset* shows the static non-linearity mapping 2 *θ* to *v*_*i*1_.

### Angular speed threshold model

Like the constant angle predicted at peak time by the *κ* function, the function

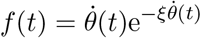

results in a constant angular velocity at peak time, 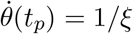. Solving for the peak time using eq. 5 yields

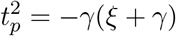

from which we deduce that the following condition on *ξ* must hold: > –*γ* > 0 (given that 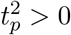). Solving for *t_p_* yields the relation

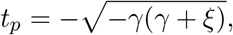

where the leading sign is selected by our use of negative times prior to collision. The square root dependence is expected from [30]. We will not consider this model further as it has not been documented experimentally.

## 5 Non-Zero Acceleration Stimuli

As explained in the previous section, the dynamics of the *η, κ*, and GF models for looming stimuli are indistinguishible modulo a static non-linearity (eq. 7), even though their functional forms differ to match the biophysics of the neurons they describe. In particular, when the models’ parameters are appropriately matched, they will predict the same angular threshold size for the neuronal peak firing rate (or membrane potential) in response to looming stimuli. This raises the question of whether it is possible to design stimuli approaching on a collision course that predict different peak response times across the models. We describe here one such type of stimuli that we call non-zero (constant) acceleration stimuli (NZAs).

### Approach with constant acceleration

If an object approaches with non-zero, constant acceleration its normalized distance is described by

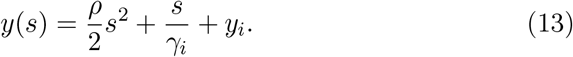

In this equation, *ρ* = *a/l* is the normalized acceleration in units of 1/time^2^, and 1/*γ_i_* is the initial normalized speed (in units of 1/time). Note that since *y*(*s*) > 0 decreases towards collision, *ρ* < 0 represents an accelerating stimulus (because the normalized distance decreases faster than for *ρ* = 0), and vice-versa for *ρ* > 0.

To compare neural model responses to an accelerating and a looming stimulus with the same projected collision time, *s_c_* = −*γ_c_y_i_*, we need to select *ρ* such that

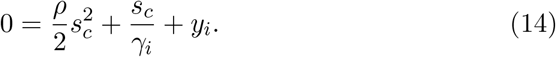

Solving for *ρ* yields

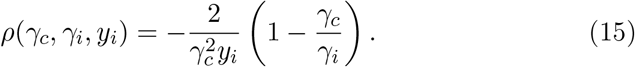

As expected, eq. 15 shows that *ρ* < 0 (acceleration) if *γ_i_* < *γ_c_* < 0 resulting in an earlier collision time than that of the looming stimulus with initial value *γ_i_*. Conversely, *ρ* > 0 (deceleration) if *γ_c_* < *γ_i_* < 0. Further, eq. 15 implies that the collision time can be made arbitrarily close to *s* = 0 (by enforcing *γ_c_* → 0), leading to *ρ* → –∞. In contrast, there is a maximal deceleration value *ρ_M_* > 0 that still leads to collision (*s* = 0). It is obtained by setting the discriminant of the quadratic eq. 14 to zero, so that its two roots coincide. This yields 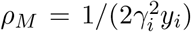 and the corresponding collision time (i.e., the double root of eq. 14) is given by *s_c_* = −1/(*γ_i_ρ_M_*) = −2*γ_i_ρ_i_*. Thus, the associated constant speed looming stimulus trajectory has *γ_c_* = 2*γ_i_*. In other words, a decelerating NZA stimulus cannot have an initial γi value smaller than half that of the looming stimulus with the same collision time.

### Examples

Fig. **5A** illustrates two NZAs with *γ_i_* = −50 ms and *γ_c_* = −20, and −80 ms, respectively. The corresponding normalized accelerations are equal to −39.3 s^-2^ and 2.45 s^-2^, respectively. Note that the first trajectory is part of an inverted parabola, while the second one is part of an upright one. In both cases, the collision time *s_c_* is one of the roots of the quadratic polynomial of eq. 13.

**Figure 5:**
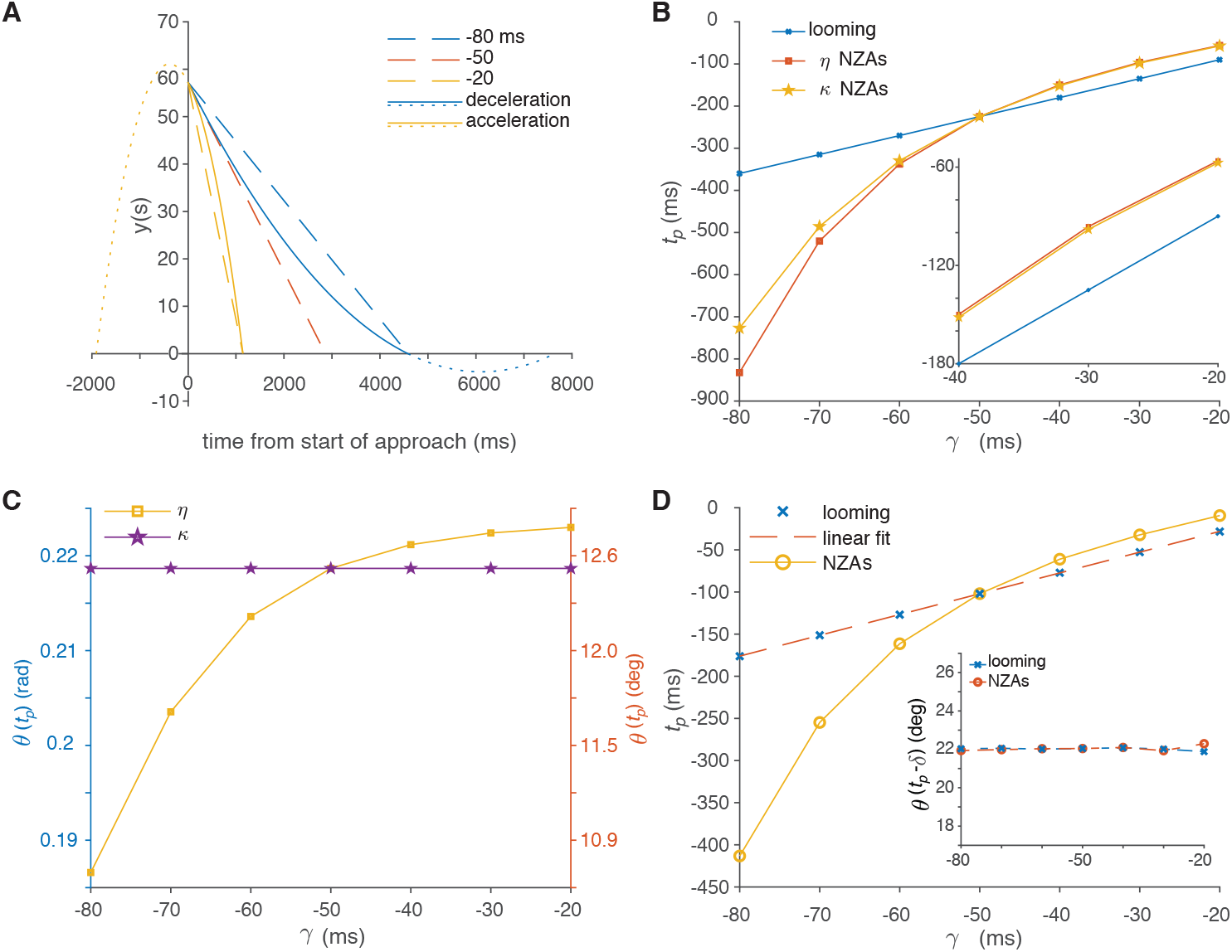
**A**, The dashed lines depict looming stimuli with three *γ* values (−80, −50 and −20 ms). The upright parabola (solid and dotted blue continuation line) represents a decelerating stimulus with the same initial *γ* value (slope) as the looming stimulus represented by the dashed red line, and the same time to collision as the looming stimulus represented by the dashed blue line. The inverted parabola represents a similar, accelerating stimulus (solid and dotted yellow line). **B**, Peak times relative to collision, *t_p_*, predicted by the *η* and *κ* models as a function of *γ* for looming stimuli and for NZAs (same parameters as in Fig. 2). The *inset* magnifies the upper right part of the graph. **C**, Corresponding peak threshold angles for NZAs predicted by the *η* and *κ* models. The latter ones are also the threshold angles for looming stimuli irrespective of model type. The left and right vertical axes give angle values in radians and degrees, respectively. **D**, Peak times of the GF model as a function of *γ* for looming stimuli and NZAs. The linear fit of looming peak times has a slope equal to *α*/2 = 2.47 and an intercept *δ* = 21.3 ms (see eq. 3). The *inset* show the threshold angle for both stimulus types as a function of *γ*.

### Comparison with insect free flight data

Although NZAs have not yet been systematically used experimentally (but see [31]), acceleration plays a role in the capture success of vinegar flies by damselflies [33]. In this work, acceleration leading to successful capture is −1.2 mm/s^2^ (using our sign convention) for *γ* = −15 ms (**Fig. 6b** of [33]). In **Fig. 6c** of the same article, deceleration leading to unsuccessful capture is equal to 0.6 mm/s^2^ for *γ* = −27 ms. Overall, the range of accelerations reported in **Fig. 6e** of [33] ranges from −3 mm/s^2^ to +1 mm/s^2^ (note that a scaling factor 10^-3^ is missing for the acceleration values in the three panels of **Fig. 6**). Taking into account the half-size of damselflies, 1.4 mm [33], this corresponds to normalized acceleration values between *ρ* = −2.14 and 0.71 1/s^2^.

**Figure 6:**
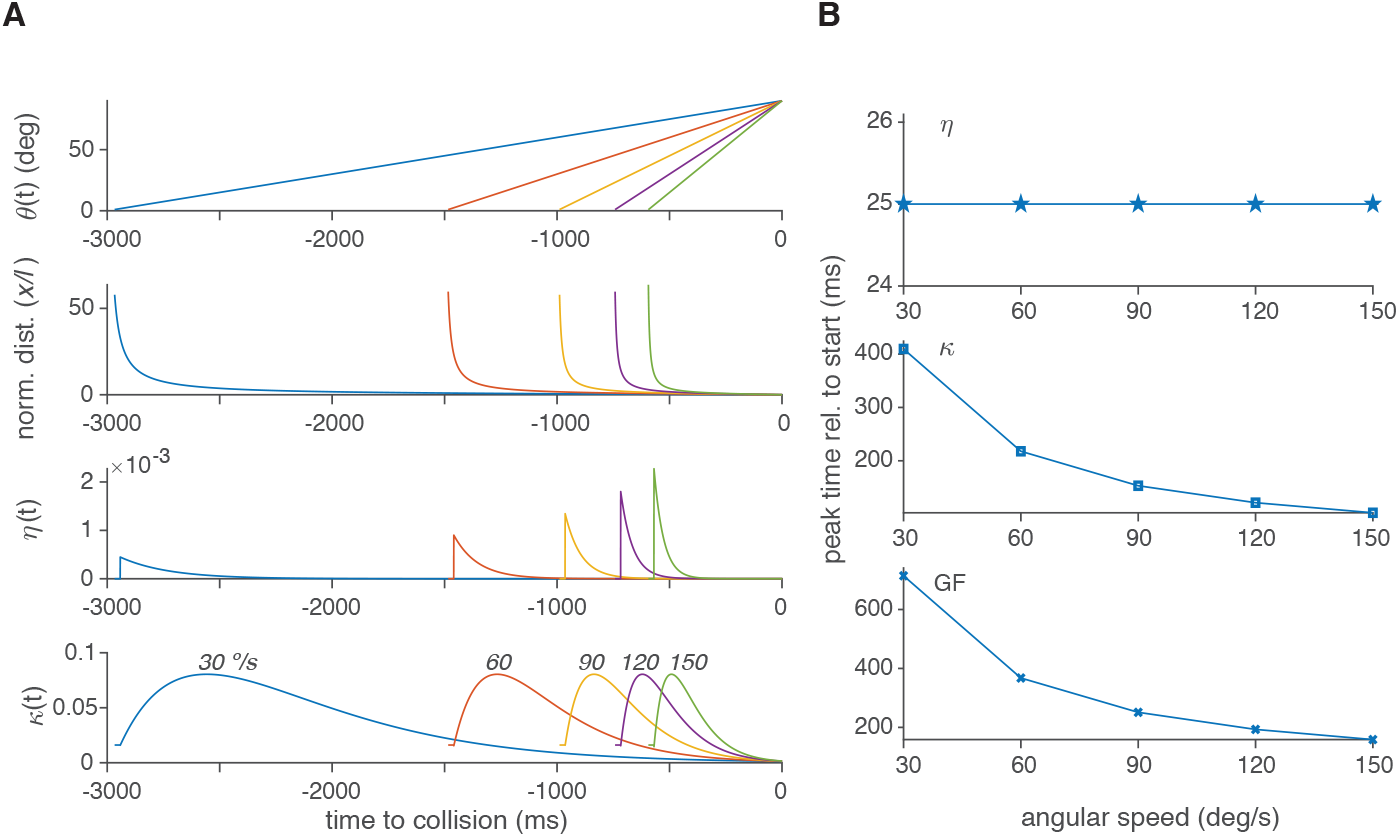
**A**, *From top to bottom*: Angular size of CAVs, corresponding normalized distance, and responses of the *η* and *κ* models for constant angular approach speeds of 30, 60, 90, 120 and 150°/s, respectively (labeled in the bottom panel). **B**, Peak times relative to start of stimulus predicted by the *η*, *κ* and GF models as a function of constant angular speed.

### Normalized distance factorization

By construction, one root of the quadratic polynomial in eq. 13 is the prescribed time to collision, *s_c_* = –*γ_c_y_i_*. The second root, obtained using the solution formula for quadratic equations, is given by

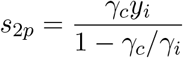

(after using eq. 15). It corresponds to the positive square root of the associated discriminant (hence the subscript ‘2p’, for ‘2^*nd*^ root, positive’). Note that if *γ_i_* < *γ_c_* < 0 (acceleration) then *s*_2*p*_ < 0. If 2*γ_i_* ≤ *γ_c_* < *γ_i_* < 0, then *s*_2*p*_ > *s_c_*, as expected (see Fig. 5**A**). Thus,

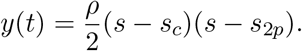

Using this factorization, we may rewrite eq. 13 in terms of time to collision,

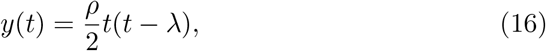

where

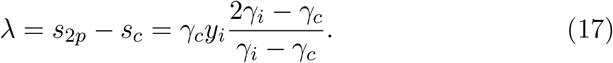

This factorization allows to determine analytically the peak times of the *κ* and *η* models.

### Peak time of the kappa model

According to eq. 8, *θ*(*t_p_*) = 1/*β* and since tan *θ*(*t*) = 1/*y*(*t*) (eq. 1), the peak time is determined by

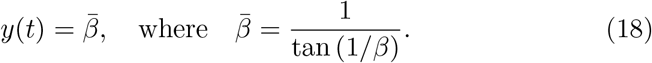

Using eq. 16 we obtain the equivalent condition for stimuli with non-zero acceleration:

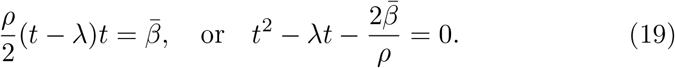

The two possible solutions are

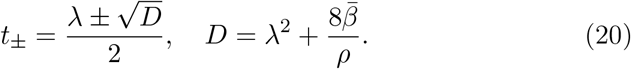

If *ρ* is positive (deceleration), *y*(*t*) is an upright parabola and its first crossing of the line 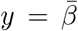 is the correct solution. Since λ = *s*_2*p*_ – *s_c_* is positive, 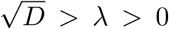 and *t*_−_ is the correct solution. If *ρ* is negative, *y*(*t*) is an inverted parabola and its second (rightmost) crossing with the line *y* = *β* is the correct solution. Since λ < 0, 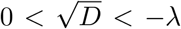 and *t*_+_ is the correct solution. Multiplying both the numerator and the denominator of either solution by *ρ*/2 and keeping track of the signs yields the unified formula,

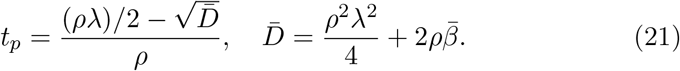

### Peak time of the eta model

The peak time is determined by eq. 2. In addition to eq. 5, we need

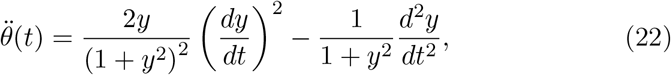

as well as

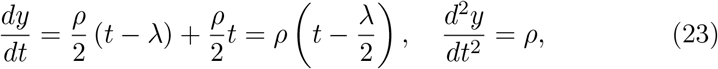

and

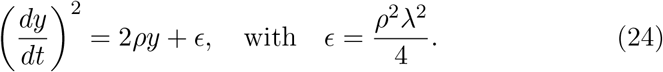

Using these results in eq. 2 leads to

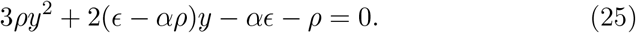

Note that if *ρ* = 0 this equation reduces to *y* = *α*/2 as expected for looming stimuli (see eq. 3).

For *ρ* > 0 the function *g*(*y*) = 3*ρy*^2^+ 2(*ε* –*αρ*)*y* –*αρ* –*ρ* is an upright parabola. Its value at *ρ* = 0 is –(*αε* +*ρ*) < 0 since *α* > 0, *ε* > 0 and *ρ* > 0. Hence, it has one positive and one negative root determined by the equations,

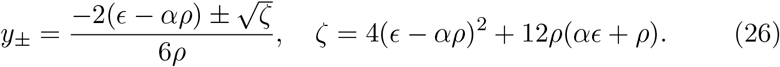

Clearly, *y*_+_ is the positive root since |2(*ε* –*α* ρ)|.

Conversely, if *ρ* < 0 then *g*(*y*) will be an inverted parabola. As shown below, under our assumptions both

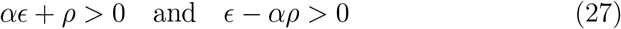

hold. The first inequality implies that *g*(*y*) has two positive roots. Combined with the second inequality this also implies that 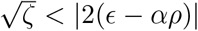 and thus *y*_+_ is the smallest root.

To obtain time relative to collision at the peak, we can use again eq. 20 but with 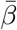 replaced by *y*_+_. Hence, the solution is

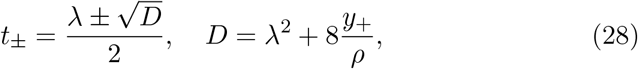

where the negative sign is selected for *ρ* > 0 and the positive sign for *ρ* < 0. Alternatively, we can use the value of tp computed from eq. 21 with the same substitution as above.

### Examples

Fig. **5B** and **C** plot the peak times and peak threshold angles predicted by the *η* and *κ* models and computed from eq. 21 using the same parameters as in Fig. 2. For decelerating NZAs (*γ* < −50 ms), both the *η* and *κ* models predict peak times occuring before those of looming stimuli with the same time to collision. Conversely, accelerating NZAs result in peak times after those of their corresponding looming stimuli. The timing difference is smallest for fastest accelerating stimuli and becomes increasingly large as *γ* becomes more negative (i.e., for larger deceleration). Accordingly, the *η* model predict increasingly smaller threshold angles than the *κ* model as deceleration increases.

### Peak time of the GF model

The peak times of the GF model were computed numerically for looming stimuli and NZAs (Fig. 5**D**). As for the *η* and *κ* models, the peak times of decelerating NZAs occur earlier than those of their associated looming stimuli, while those of accelerating NZAs occur later. A linear fit of looming stimuli peak times allows to compute the associated threshold angles using eqs.3 and 4 (Fig. 5**D**, *inset*). This reveals that the threshold angles of the GF model are nearly constant for NZAs. Thus, the GF model resembles more closely the *κ* than the *η* model in this respect.

### Proof of the inequalities in eq. 27

To prove the first inequality, note that when *ρ* < 0 we have *γ_i_* < *γ_c_* < 0 and hence 0 < *γ_c_*/*γ_i_* < 1. Setting *x* = 1 – *γ_c_*/*γ_i_* this implies 0 < *x* < 1. Using eqs.15 and 17, the first inequality may rewritten as

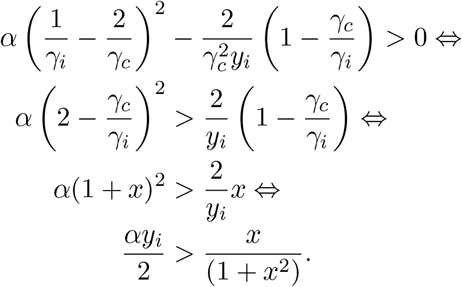

As the maximum value of the right hand side over [0; 1] is 1/4, we obtain *αy_i_* > 1/2, which will always be satisfied under our typical conditions (e.g., *α* = 9 and *y_i_* = 76.4, see Figs. 1 and 2).

Consider now the second inequality, which may be rewritten as

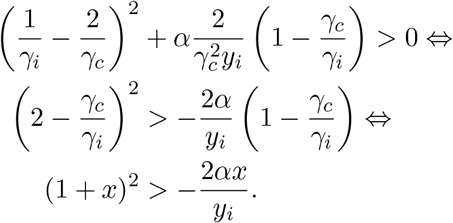

This last inequality will always be satisfied since the right hand side is smaller than 0.

### Testing model predictions on surrogate data

Although the *η* and *κ* (or GF) models considered above make different predictions for the peak times and angular threshold values associated with NZAs, it remains to be seen whether the stochastic variability of neuronal responses allows to distinguish them experimentally. This question can be addressed theoretically for the LGMD/DCMD neurons since the experimental variability of their peak responses has been characterized for looming stimuli [15]. If we assume this variability to be normally distributed [15], it can be extrapolated to NZAs under the assumptions of the *η* or *κ* models. We can then test the ability to detect a fixed **κ** or variable (*η*) angular threshold for NZAs using surrogate data generated from these variability models.

### Surrogate data for the kappa model

For the *κ* model, we assume that the fixed threshold angle takes the value *θ_t_* = 12.5° as is typical for the LGMD/DCMD neuron [15]. In addition, we assume a fixed angular error, which we deduce from Fig. 7 of [15] to be equal to *σ_θ_t__* = 3.1°/2 = 1.55° (note the halving since the variability given there is for the full angle, *θ_f_*). Under these assumptions, the angular threshold variability comes from a normal distribution with mean *θ_t_* and variance 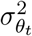,

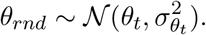

Further, we assume a typical delay between peak time and angular threshold, *δ* = 25 ms [15]. Using eqs.18 and 3, the corresponding peak random times for looming stimuli are given by

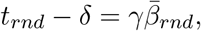

where 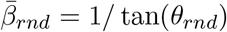. According to eq. 19, the peak random times for NZAs are given by

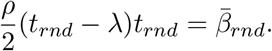

The value of *t_rnd_* is obtained from 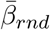 using eq. 20.

### Surrogate data for the eta model

According to eq. 2, the *η* model is characterized by a relative angular acceleration threshold, 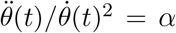. For looming stimuli, we deduce from eqs. 5 and 22 that,

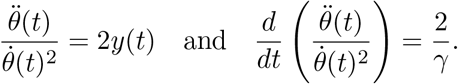

Hence, to first order,

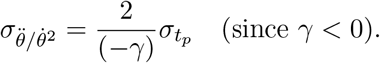

Comparing with eqs. (8) and (9) of [15] we deduce that

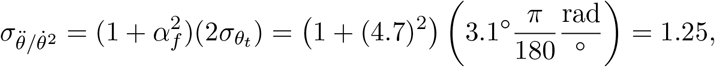

where the value of *α_f_* is taken from Fig. 4 of [15].

To generate random peak times of the *η* model, we start with a random variable representing the relative angular acceleration threshold,

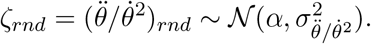

For a looming stimulus, *ζ* = 2*y*(*t* – *δ*) which leads to 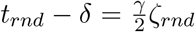. For NZAs, eqs.5 and 22 lead to

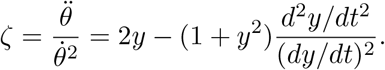

Using eqs.23 and 24, we obtain

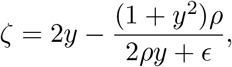

which leads to

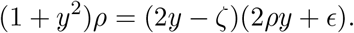

This last equation is identical to eq. 25 and, for a given value of *ζ_rnd_*, it is solved as in eq. 26 to obtain *y*_+_(*ζ_rnd_*). We can then use this value in eq. 28 to compute *t_rnd_* – *δ*.

### Simulation results

Using the two surrogate data models described above, we simulated 10,000 synthetic data sets consisting of 10 peak times for looming stimuli with *γ* values equal to −20, −30, −40, −50, −60, −70 and −80 ms and 10 peak times for NZAs with the same γc values. The peak times of looming stimuli were fitted with a linear model as in [15] to determine an estimated slope and delay (see eq. 3). The estimated delay was then used to compute the threshold angles, *θ*(*t_p_* – *δ*), using the synthetic *t_p_* values for the NZAs. Since the computed threshold angles are non-linear functions of *θ_rnd_* and *ζ_rnd_*, they may not follow a normal distribution. Hence, we tested whether they were identical across *γ* values using the Kruskal-Wallis procedure, a non-parametric analysis of variance.

For the *η* model, 81 percent of synthetic data sets had threshold angles that were significantly different across *γ* values. In contrast, for the *κ* model only 22 percent of synthetic data sets had significantly different threshold angles. These results suggest that LGMD/DCMD data for paired looming stimuli and NZAs might be able to distinguish the two models despite its expected experimental variability.

Further, based on an Anderson-Darling test 38 percent of synthetic *η* data sets had computed angular threshold values that were not normally distributed. For the *κ* model, only 7 percent of synthetic data sets were not normally distributed. These results justify the use of the non-parameteric Kruskal-Wallis test.

## 6 Constant Angular Velocity Stimuli (CAVs)

A second type of stimulus with varying acceleration leads to different responses in the *η, κ* and GF models. This stimulus type simulates objects approaching at a constant angular velocity. If *θ_i_* is the initial and with *θ_c_* the final half-angle at collision (in our case 1° and 90°, respectively), the trajectory is described by the equation

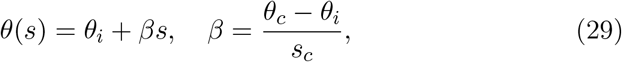

where *s* is time from stimulus start, *s_c_* is the collision time and *β* is the angular speed (in units of rad/ms). If time is computed relative to collision, *t* = *s* - *s_c_*, then substituting *s* = *t* + *s_c_* in the right hand side of eq. 29 yields

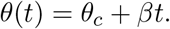

The corresponding normalized distance is obtained from eq. 1 and is illustrated for a set of constant angular velocities in Fig. 6**A**, immediately under the corresponding constant angular velocity traces (second panel from the top). The selected angular velocities span the range observed at the peak firing rate time of the LGMD neuron for looming stimuli between −80 and −20 ms. As can be seen from this figure, CAVs correspond to an extreme case of deceleration around collision time. The instantaneous normalized speed of approach is given by

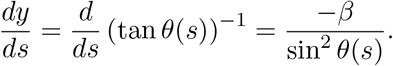

Hence, the normalized speed is largest at the beginning of the approach, when *θ_i_* = 1°, and smallest at the end of approach. In general, the smaller the initial angle the larger the initial speed since sin *θ* → 0 as *θ* → 0, while at collision time *dy*/*ds* = −*β* since sin *θ* = 1 when *θ* = 90°.

The response of the *η* and *κ* models are depicted in Fig. **6A** (bottom two panels, respectively). The *η* function predicts an initial transient to the decelerating stimuli locked to stimulus motion onset, while the *κ* model predicts a peak at its preferred angular size. Correspondingly, the peak time of the *η* model follows with a fixed delay (*δ*) the stimulus onset, whereas the delay in response of the *κ* model decreases with increasing angular velocity (Fig. 6**B**). The GF model predicts a peak response time relative to stimulus onset that is similar to that of the *κ* model.

## 7 Discussion

In this work, we investigated three models describing the responses of looming-sensitive neurons in grasshoppers, vinegar flies, and goldfish, respectively. When presented with looming stimuli, the three models predict a fixed threshold angular size at the time of peak neural response, independent of the stimulus parameter *γ* (modulo a delay). In contrast, we show that for stimuli with accelerating trajectories the three models predict different peak response times and threshold angles. Our results clarify the relation between the models and provide new tools to study and distinguish them experimentally.

The *η, κ* and GF models were formulated using different experimental constraints and biophysical considerations [16, 1, 34, 24]. The *η* model of grasshoppers predicts an increase in peak firing rate as *γ* increases (e.g., from −80 to −20 ms; Fig. 2**A**). Experiments confirmed this dependence for the peak LGMD/DCMD firing rate, although it is not as strong as predicted [15]. Similarly, combined behavioral and electrophysiological data suggest that a fixed number of spikes is required to generate jump escape behaviors, though again not from stimulus start as predicted [13]. In contrast to the *γ* dependence of the peak *η* model firing rate, the *κ* model predicts a fixed peak response (Fig. 2**B**). This is what was observed in intracellular recordings of the goldfish Mauthner cell [24]. In the GF model of *Drosophila melanogaster*, the peak response to looming stimuli is also predicted to be *γ* dependent, though not as strongly as in the *η* model (Fig. 4).

In grasshoppers, the variability of LGMD/DCMD firing rate responses to looming stimuli across animals has been studied over a broad range of *γ* values [15]. This showed a large variability across animals, requiring a generalization of the *η* model that adds a static non-linearity to accommodate the data (see eq. 7; [15]). This generalized model is sufficiently flexible to also describe the GF or the *κ* model. We illustrated this by providing a closed (and invertible) formula mapping the *κ* model onto the *η* model (see eq. 11 and Fig. 3). This observation raised the question of whether other stimuli might better differentiate the *η*, *κ* and GF models.

We took a hint from work on the prey capture of vinegar flies by dam-selflies, showing that predator acceleration is ethologically relevant [33]. Thus, our first set of stimuli adds a non-zero constant acceleration term to looming stimuli. The normalized acceleration values of these non-zero acceleration stimuli encompass the range relevant for prey capture by damselflies. And indeed, we found that NZAs lead to different model responses, with the *κ* and GF models predicting a constant angular threshold while the *η* model does not (Fig. 5). The reason is that the *η* model is not a ‘true’ angular size threshold model, but rather a relative angular acceleration threshold model (eq. 2). These two threshold models (angular size and relative angular acceleration) happen to make identical predictions for looming stimuli. Notably, the sensitivity to angular acceleration of the LGMD/DCMD neurons has been documented when studying their responses to pseudo-naturalistic stimuli [25, 29].

A second hint came from early DCMD recordings to constant angular velocity stimuli, which showed a fixed delay of peak response following the onset of stimulation [16]. In contrast, our results show that both the GF and *κ* model predict a decreasing delay with an angular speed increase (Fig. 6**B**). CAVs exhibit strong deceleration, and accordingly the differences between the models are stronger than for NZAs. Yet, although NZAs produce more subtle changes across models, surrogate data based on variability extrapolated from neural DCMD responses to looming stimuli suggest that they might still be detectable experimentally.

A related question is whether the different model predictions could be tied to behavioral escape responses for accelerating stimuli as already shown for looming stimuli (e.g., [13, 11, 33, 3]). Because behavioral variability is usually considerably higher than neuronal response variability [13], this is expected to be more difficult. For GF-mediated behavioral responses an additional complication stems from the fact that the probability of response starts low and increases with *γ*, which is also when the differences in model predictions are the smallest [33].

In summary, this works brings a better understanding of the relationship between neural models of collision detection. Further, it also establishes the groundwork to compare experimentally their predictions for accelerating stimuli, a topic pursued in a companion paper.

## Statements and Declarations

### Data availability statement

The Matlab code required to reproduce all the figures and the statistical analysis of surrogate data for non-zero acceleration stimuli presented in sec. 5 is available online at Mendeley Data (repository DOI).

### Author contributions

FG, TP and RD designed the work; FG carried out the mathematical derivations for model predictions and wrote the computer code to analyze the results; FG prepared the figures; FG wrote the manuscript with input from TP and RD.

All authors approved the final version of the manuscript. All persons designated as authors qualify for authorship, and all those who qualify for authorship are listed.

## Acknowledgments

We would like to thank Dr. Gwyneth Card for sharing the data needed to compare NZAs with the acceleration values observed in damselflies during naturalistic prey capture experiments.

